# No Evidence for a Vestibular Contribution to Object Motion Prediction

**DOI:** 10.64898/2026.02.04.703809

**Authors:** Björn Jörges, Laurence R. Harris

## Abstract

Humans can predict an object’s motion better if its movements are consistent with gravity. Here we investigate whether this may be due to an internalized strong Earth gravity prior or to vestibular cues reporting instantaneous information about gravity. These two directions can be separated using virtual reality by providing strong visual cues to the direction of up which may or may not be aligned with true gravity. Participants were presented with a ball travelling on a parabola path simulated with either downward acceleration created by simulated Earth’s gravity (1g) or inverted gravity (-1g) resulting in the ball curving upwards. In both types of trial, the ball disappeared at between 57.5% and 75% of its full trajectory – after it had started its descent in the case of 1g or ascent in the case of -1g. Participants pressed a mouse button when they judged the ball to have got back to the height at which it was launched. Participants were either standing or supine. There were no differences in the estimated time to reach the indicated level between the 1g and -1g simulations nor between lying and sitting.A control experiment showed that reaction times were also not significantly different while lying supine versus while upright. Fiinally, to isolate a vestibular component we compared performance during disruptive galvanic vestibular stimulation (dGVS) or during sham stimulation. As when lying supine, the perceived time for the ball to reach the target height was not significantly different between dGVS and sham stimulation. Overall, participants were – surprisingly – no better at anticipating 1g motion compared to -1g motion, and we also don’t find a significantly influence of the vestibular system more generally.

## Introduction

Over the past two decades, a growing consensus has emerged that humans use internalized knowledge of Earth gravity to predict the motion of objects in their environment (Baurès & Hecht, 2011; Bosco et al., 2015; Ceccarelli et al., 2018; De Sá Teixeira, 2014, 2016; Freitas & De Sá Teixeira, 2021; Gaveau et al., 2011; Indovina et al., 2005; Jörges et al., 2018; Jörges & López-Moliner, 2017, 2019, 2020; Velado et al., 2024; Zago et al., 2004b, 2004a, 2008, 2009, 2010, 2011a). Paradigmatically, participants are shown objects that either accelerate downwards (at 1g) or upwards (at - 1g). The object then disappears, and participants predict when the now-invisible object reaches a certain landmark. Typically, participants overestimate the time it takes the object to reach the landmark under -1g conditions in comparison to 1g trials. The underlying mechanism is thought to be that the human visual system is highly insensitive to arbitrary accelerations and therefore tends to neglect them in motion extrapolation (Benguigui et al., 2003; Bennett & Benguigui, 2013). However, when the acceleration acting upon an object corresponds to Earth gravity, the perceptual system seems able to compensate for this shortcoming by using internalized knowledge of Earth gravity. This allows humans to predict the motion of objects more accurately under 1g conditions.

Based on the experiments conducted so far, it is impossible to ascertain whether these performance benefits for 1g object motion are (a) due to a strong Earth gravity prior or (b) due to ongoing vestibular signals indicating that the observer is in a 1g environment and should expect external objects to behave accordingly. The conceptual difference between the two is that the strong Earth gravity prior has been built up over a lifetime interacting with objects moving in accordance with Earth gravity. Some evidence even points to such a prior being innate (Bardi et al., 2014; Vallortigara & Regolin, 2006) or being acquired early in development (Bertenthal et al., 1987). Experimental evidence has shown that this prior is very robust even in the presence of feedback in environments with changed visual gravity (McIntyre et al., 2001; Zago et al., 2004b). The prior is activated in appropriate situations, e.g., when viewing object motion that is embedded in a pictorial environment (Zago et al., 2011b) when it can aid in the prediction of object motion in situations where this might be necessary or useful, such as when anticipating motion during an occlusion or to compensate for neural delays (Zago et al., 2008).

In contrast to accessing a gravity prior, ongoing vestibular signals could be used to keep track of the direction of gravity pursuant to the observer’s movements and their posture in their environment (Kobel et al., 2021; Merfeld et al., 1999). The perceptuo-motor system could then potentially access this representation of gravity to aid in predicting motion. Figure 1A and B illustrate these potential contributions: for an upright observer, both the gravity prior (red arrow) and the vestibular input (blue arrow) point in the same direction. Both could serve to help compensate for the inability of the perceptual system to extract and use the acceleration acting on the tennis ball based purely on online visual cues (green arrow).

**Figure 1:**
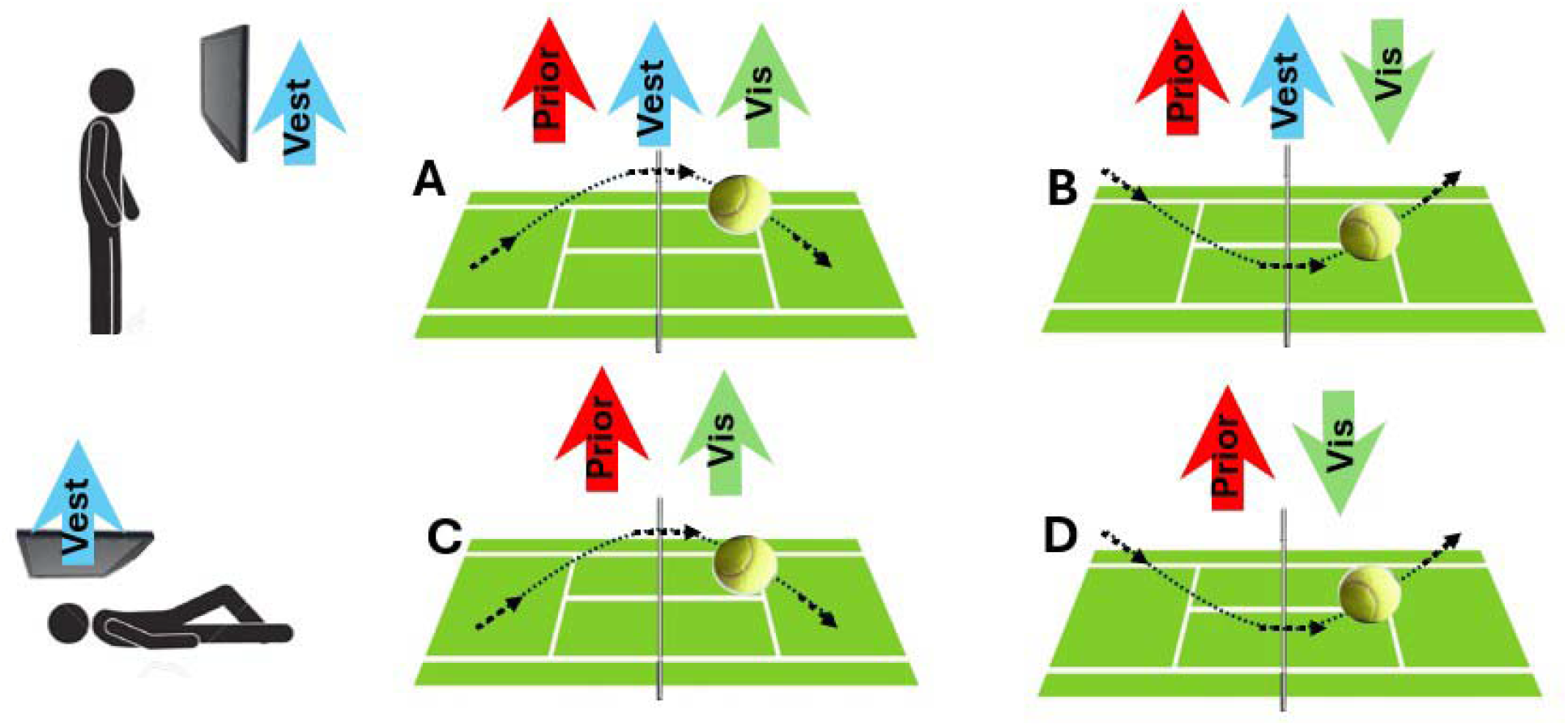
Diagram detailing the different scenarios covered by our hypothesis. The arrows represent the direction of “UP” signalled by each means (red= gravity prior aligned with the scene, blue = gravity as determined by vestibular processing, green = visually simulated acceleration of the object). In A and B, the gravity prior and vestibular input are aligned. In C and D, the observer is lying supine while the visual scene remains aligned with their body, i.e., “down” in the scene is towards their feet. When lying, vestibular input is orthogonal to the visual scene and therefore to the gravity prior, which makes it task-irrelevant. In A and C, object motion unfolds according to Earth gravity, while B and D are scenarios where the object obeys a simulated inverted Earth gravity, i.e., it accelerates upwards rather than downwards.

Several studies have addressed the interplay between vestibular signalling and the gravity prior, e.g., for the perception of visual acceleration (Miwa et al., 2019), the duration of vestibular and/or visual motion intervals (Delle Monache et al., 2025) and in the context of representational momentum and gravity (Freitas & De Sá Teixeira, 2021). However, to our knowledge only two experiments have tried to assess the relative contributions of the strong gravity prior and ongoing gravity signalling directly in the context of object motion *prediction*. McIntyre et al. (2001) conducted an experiment with astronauts and compared their performance catching downwards moving balls on Earth with their performance onboard the International Space Station (ISS). Measuring muscle activation, they showed that astronauts initiated their responses too early even after 15 days in space, indicating that they expected the downwards moving object to accelerate (rather than to travel at a constant speed as it actually does in microgravity). A shortcoming of this study was the unavoidable downside of using real-world objects: McIntyre et al. used actual balls, which were associated with accelerating visual motion on Earth but motion at a constant speed in microgravity, rather than comparing the same (constant speed) type of motion in space and on Earth. While the authors argue (convincingly, in our opinion) that evidence from muscle activation onset is sufficient to support the internal representation hypothesis, the stimulus was not fully controlled. Baurès and Hecht (2011) investigated in a VR study on Earth whether the motion of an object moving towards an observer was predicted differently when participants were lying supine compared to when they were lying prone. While supine, participants should have expected the object’s to accelerated with Earth gravity, while it would have accelerated against Earth gravity (i.e., at -1g) when participants were prone. The authors found differences in line with a 1g expectation but only at the longest occlusions (>= 2.5s). Importantly, a rudimentary visual environment (a brick-walled tunnel) was used and the stimulus, while visible, was moving at a constant speed: both of these factors may have impacted the ability of the perceptual system to recruit the strong Earth gravity prior. The lack of an acceleration control in the McIntyre study (McIntyre et al., 2001) and uncertainty regarding the activation of the gravity prior in the Baurès and Hecht study (Baurès & Hecht, 2011) mean that determining whether the strong Earth gravity prior or ongoing vestibular signalling are responsible for performance benefits for 1g targets over -1g targets is not yet fully resolved.

The present study addresses the relative contribution of the strong prior versus ongoing vestibular signalling in a novel way: if vestibular signalling were responsible for the differences between 1g and -1g motion reported in the literature and reviewed above, then making the vestibular input irrelevant should make any difference between 1g and -1g (Fig 1a and b compared to Fig 1 c and d) disappear. If, conversely, the difference between 1g and -1g were maintained under such circumstances, this would mean that time-to-contact estimation relies predominantly on the strong Earth gravity prior. In our study, we made the vestibular input task-irrelevant by tilting observers into a supine position while keeping the visual stimulus aligned with their bodies. That is, visually, the orientation of the virtual environment indicated that gravity was aligned with their bodies, while vestibular cues indicated gravity at 90° from this. This is illustrated in Figure 1C and D where the vestibular input (blue arrow) is orthogonal to the visual scene which serves as a reference frame for the strong gravity prior (red arrow). This makes the vestibular component task-irrelevant and removes it as a potential cue to help predict object motion. We hypothesized that the contribution of the prior will decrease when observers lie supine (where visual, vestibular and somatosensory cues are put into conflict) or in the presence of disruptive galvanic stimulation.

## Methods – Experiment 1: Sitting versus Lying Supine

### Overview

Here, we compared how participants judged the movement of a free-flying ball in the presence of conflicting information about the direction of gravity compared to their performance with coherent information. When lying supine, the visual system (fooled using virtual reality) told them up was aligned with their body, but the vestibular and somatosensory systems told them it was orthogonal to that direction.

### Participants

26 participants performed the task. One participant (n = 1) did not finish the experiment due to dizziness and was therefore excluded. All participants were recruited from the York University undergraduate research participant pool and received course credit for their participation. They all had normal or corrected-to-normal eyesight and gave their written informed consent prior to the experiment. This study was conducted according to the principles of the Declaration of Helsinki. This research was approved by the York University Office for Research Ethics under the protocol numbers e2019-447 (Experiment 1 and Control Experiment) and e2024-006 (Experiment 2). All participants gave their written consent. Data collection for Experiment 1 occurred between May and July 2023.

### Apparatus

The stimulus, programmed in Unity (v2021.3.6f1), was presented using a VIVE Pro Eye head-mounted display (98° visible field-of-view vertical and horizontal, 90.46° binocular overlap, 1440 x 1600 pixel resolution per eye). Participant responded using a finger mouse either while standing upright or lying supine on a bed.

### Stimuli

Participants were immersed in a stereo rendering of a virtual office environment that provided depth cues (see Figure 2). On each trial, participants were shown a free-flying soccer ball (22cm diameter) that was fired from a cannon with a vertical velocity randomly drawn from the interval [5.5; 7 m/s] (1g motion) or the interval [5.5; 7 m/s] (-1g motion) and a horizontal velocity randomly drawn from the interval [2.5; 3.5 m/s] (left-to-right motion) or from the interval [-3.5; -2.5 m/s] (right-to-left motion). This made for initial angular velocities of 6 to 7.8 m/s and launch angles relative to the horizontal of between 57.5° and 70.3°. The ball could be affected by Earth gravity (9.81 m/s^2^) or inverted Earth gravity (-9.81 m/s^2^, i.e., an upwards acceleration). That is, the ball followed a parabolic motion profile in which it started out travelling up (1g) or down (-1g) before gravity pulled it in the opposite direction (down for 1g or up for -1g) such that, at the end of its trajectory, the ball had returned to its initial height. At the beginning of each trial, the ball (“target”) and a horizontal red bar (reaching from one wall to the other; “plank”) appeared 10 m in front of the observer. The target always began its motion precisely at the bar (shot out of a red cylinder mimicking a cannon) and its start position was adjusted for each condition such that it crossed the observer’s straight-ahead at the peak of its flight. The height at which the plank and the cylinder appeared was either 2.6 m above the simulated ground (for 1g trials) or 4.6 m above the simulated ground (for -1g trials), with a total room height of 7.2 m. The participant’s viewpoint was simulated exactly in the middle of the room and if participants looked around the virtual environment, they could see that they were standing on a platform suspended by the chains visible in Figure 2. With these parameters, trajectories were centered both vertically and horizontally in front of the observer.

**Figure 2.**
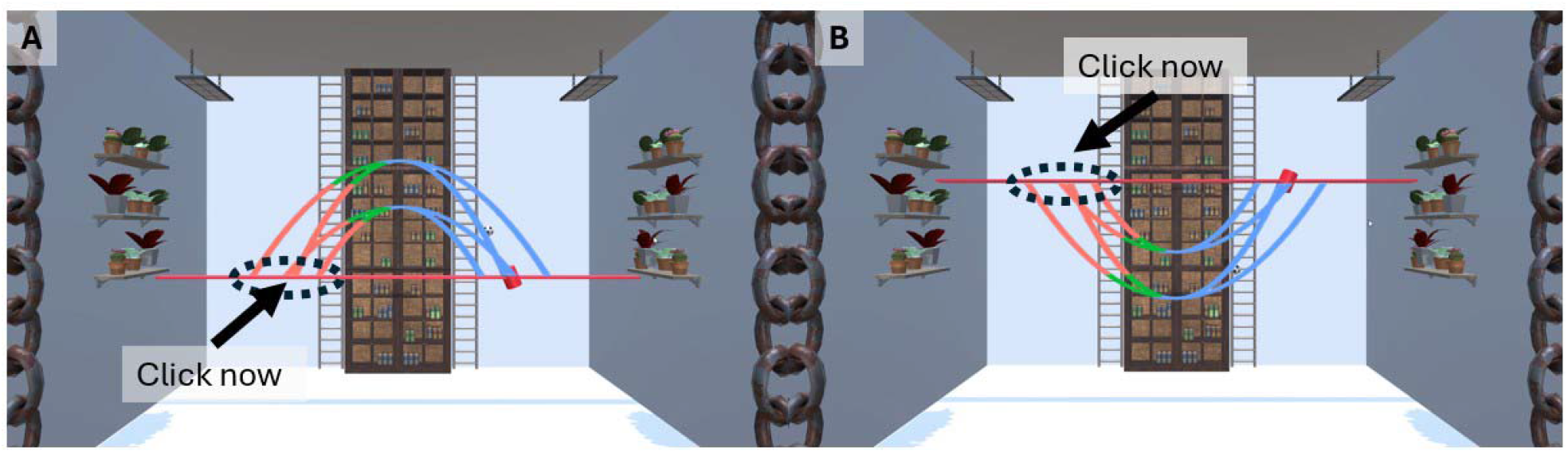
Screenshots from the experiment. Overlayed on the screenshots are the most extreme trajectories, i.e., those with initial vertical velocities of 5.5 m/s and 7 m/s and horizontal velocities of 2.5 m/s or 3.5 m/s. During the blue part of the trajectory, the object was always visible. The green parts denote the window where it could disappear (between 57.5% and 72.5% of the whole flight duration) and the object was always invisible during the red parts. The dotted black circle indicates when participants had to press the button in order to respond most accurately. **A.** 1g condition. **B**. -1g condition. Please note that the object could not only travel right to left as indicated in the figures but also left to right.

500ms after the plank and the red cylinder became visible, the target started its motion and was visible for a percentage of the total time it took for the object to complete its parabola and hit the plank (its “flight time”). This visible percentage was chosen randomly from the interval [57.5%; 72.5%], after which the ball disappeared. Participants were asked to indicate by button press the moment the now-invisible target would return to its initial height, that is, when it would hit the red plank. Occlusion durations were between 308ms and 607ms. All combinations of initial vertical velocities, horizontal velocities, occlusion durations and direction of gravity were interleaved.

Before proceeding to the main experiment, participants performed a short training of 8 trials (in whichever condition they were tested in first) where the ball reappeared upon button press, indicating how accurate their estimate had been. There was no inclusion criterion applied and participants moved to the next stage of the experiment regardless of their performance.

For the main part of the experiment, participants performed 160 trials while standing upright and 160 trials while lying supine. Including administration of instructions and set-up of each individual participant, the experiment took about 25 minutes to collect all trials in both postures. The 160 trials were composed of 80 repetitions for each gravity direction. Within each of these 80 trials, we randomized initial vertical velocities, horizontal velocities and the occlusion percentage as described above. For both postures, the relation between the participant’s body and the virtual environment was the same, i.e., the participants’ feet were oriented towards the simulated floor, and we counterbalanced the posture with which participants started (i.e., 13 participants started lying supine and 12 participants started standing upright). You can view a video of the stimulus (including the training) on Open Science Foundation (main experiment: https://osf.io/vf7qm/files/3u96x; training: https://osf.io/vf7qm/files/ca2fb).

### Data Analysis

We first computed the temporal error as the difference between the occluded flight time and the participants’ reaction time, with positive values indicating too-late responses and negative values indicating too-early responses. We excluded participant p09 because they experienced nausea in the standing posture and had to abort testing before completing the whole experimental block. We also excluded all those trials as outliers where the extrapolated flight time was higher than two times the occluded duration as trials where the participant hadn’t paid attention to the stimulus (a total of 675 out of 10240 trials or 6.6%). We then fitted a linear mixed model using the lme4 package (Bates et al., 2015) with the timing error as the dependent variable, Gravity (1g versus -1g), Posture (Standing versus Lying), as well as the initial vertical velocity and the initial horizontal velocity as fixed effects, as well as random intercepts per participants and random slopes for Gravity, Posture, vertical velocity and horizontal velocity. This model was chosen because we expected a main effect of Gravity (with later responses for -1g than for 1g) and an interaction between Gravity and Posture (with a smaller difference between 1g and -1g while supine versus while upright). For the random effects structure we follow a “keep it maximal” approach (Barr et al., 2013), and following our own observations in previous studies and simulations showing that representing variables as random effects, but not as fixed effects, we also added the vertical velocity and the horizontal velocity as fixed effects. In the Wilkison and Rogers notation, this model reads as follows:

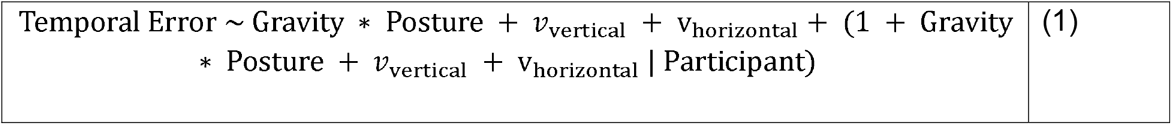

We then used the *confint()* function from base R to estimate 98.3% confidence intervals on all fixed effects to ascertain statistical significance at an alpha level of 0.05/3 = 0.017 to account for the fact that we are reporting three experiments in this paper.

## Results

### Participants are color-coded

We found no significant impact of gravity (1g versus -1g) or posture on performance. For a detailed breakdown of all difference contrasts for all fixed effects in the linear mixed model please refer to Appendix A. The results are visualized on a population level in Figure 3A and on a participant level in Figure 3B.

**Figure 3.**
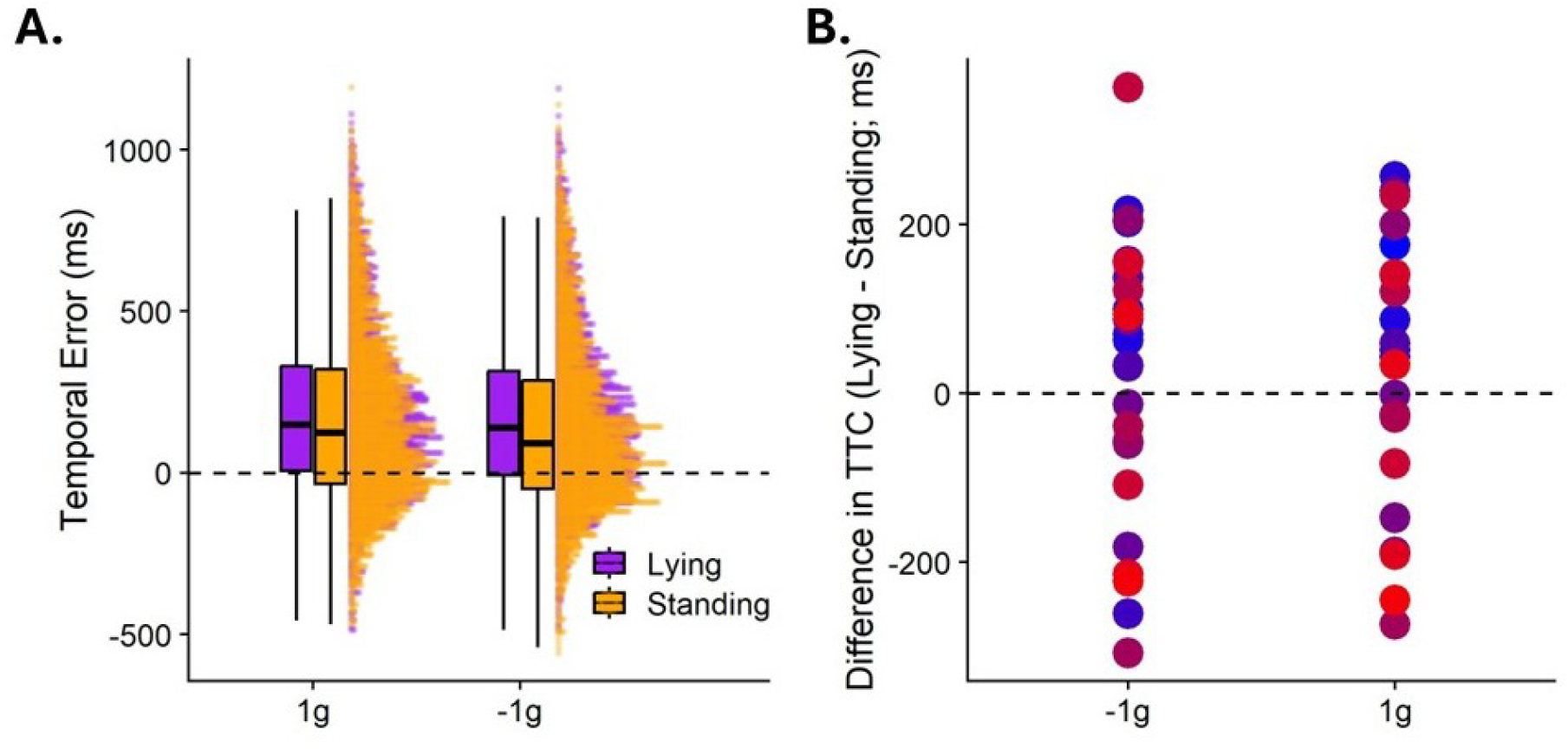
A. Full distributions of temporal errors (y axis) for each gravity level (1g and -1g, indicated on the x axis) and posture (Lying supine versus Standing upright, color-coded). The boxplots indicate the 25^th^ and 75^th^ percentile (box), while the whiskers indicate the 2.5 times the interquartile range, and the horizontal bar in the middle corresponds to the median. B. Mean difference in TTC between Lying and Standing (y axis) for the two gravity conditions (x axis). Participants are color-coded.

**Figure 4:**
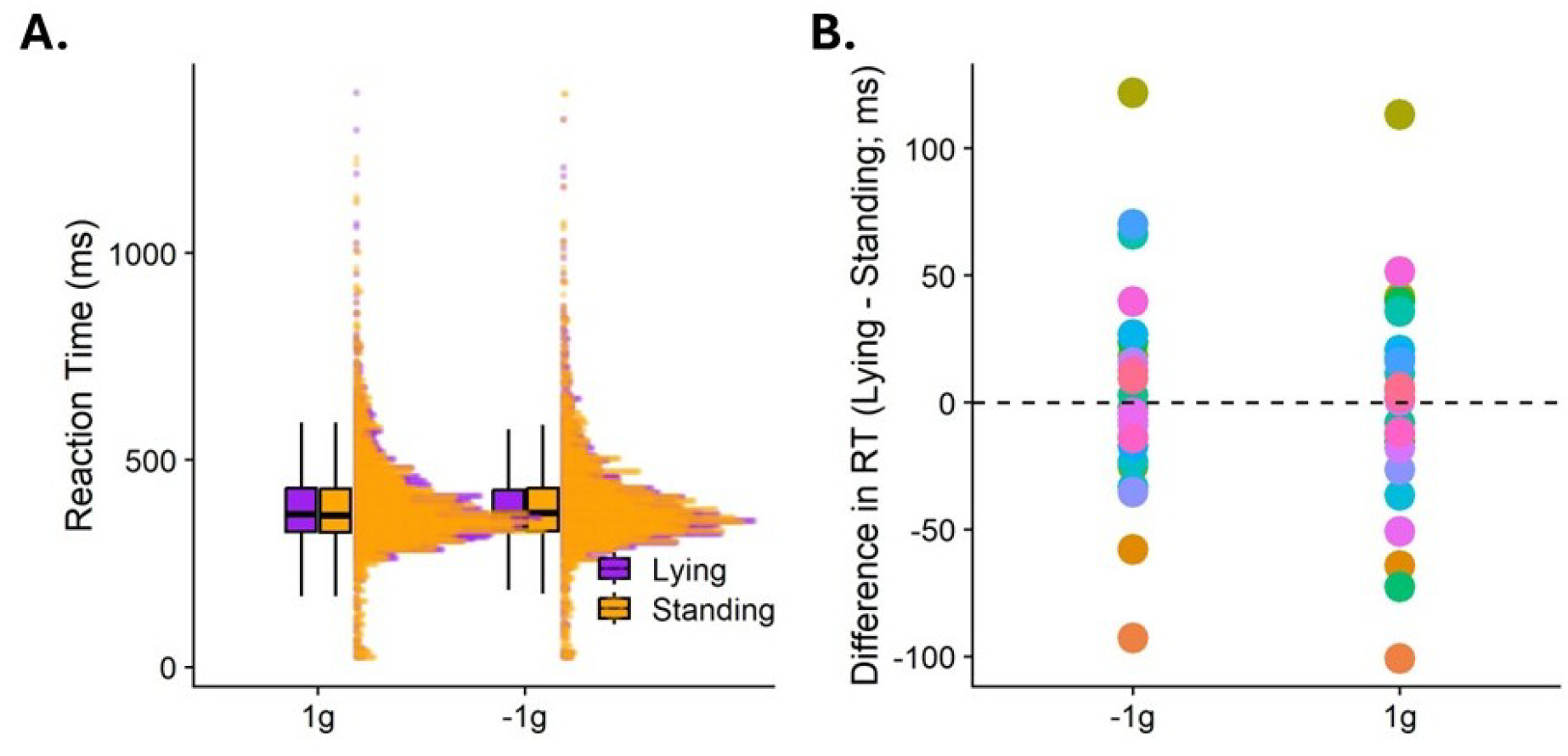
A. Full distributions of reaction times (y axis) per gravity level (1g versus -1g, x axis) and posture (Lying supine versus Standing upright, color-coded). The boxplots indicate the 25^th^ and 75^th^ percentile (box), while the whiskers indicate the 2.5 times the interquartile range, and the horizontal bar in the middle corresponds to the median. B. Mean difference in TTC between Lying and Standing (y axis) for the two gravity conditions (x axis).

## Control Experiment

To determine whether potential differences in time-to-contact were due to differences in reaction time (e.g., due to more sluggish reactions while lying down versus sitting upright), we devised a reaction time control experiment. In this experiment, the same ball appeared at the same time it would have disappeared in the main experiment (i.e., at between 57.5% and 72.5% of its trajectory) and participants had to press a button as soon as they spotted it. 26 new participants (recruited from the York University undergrad participant pool) completed 160 trials per posture (80 trials per gravity direction, within which initial vertical velocity, horizontal velocity and occlusion percentage were randomized as for Experiment 1). Data collection for this experiment occurred in February 2025. We employed the same statistical analysis as above to ascertain whether there was any population-level difference in reaction times between lying supine and standing. 35 out of 8320 total trials were excluded (0.4%).

### Participants are color-coded

**Error! Reference source not found**. 4 depicts the reaction times measured in Experiment 2. We found no significant differences between standing upright and lying supine or any of the other variables represented in the statistical model. Please see Appendix A for detailed difference contrasts and confidence intervals.

## Experiment 2: Disruptive Galvanic Vestibular Stimulation

In Experiment 1, we found no evidence that time-to-contact was overestimated when lying supine versus when standing upright. However, our research question was specifically about vestibular signalling, and there is a whole host of sensory cues that differ between lying supine and standing upright, such as blood distribution in the body, haptic cues from pressure receptors on the skin and kidneys, and even more cognitive factors such as task context (Alberts et al., 2016; Bronstein, 1999; Mittelstaedt, 1996). To further isolate whether changes in vestibular signalling specifically played a role in time-to-contact estimation we employed disruptive galvanic vestibular stimulation (dGVS) to selectively degrade the vestibular signal. dGVS massively increases the noise in the vestibular system, and if we could observe differences when comparing performance during dGVS with results obtained during sham stimulation where the current was applied not to the vestibular system but to the neck.

The vestibular stimulation consisted of a small current (50mA) applied through electrodes on the mastoid processes behind the ears. A reference electrode was placed in the center of the forehead. The electrodes were 3.25 cm diameter round carbon-conductor electrodes (9000 series electrodes; Empi et al., USA). A GVS system (Good Vibrations Inc.), generated the vestibular stimulus controlled by a PC. The vestibular stimulus was a sum-of-sines waveform with dominant frequencies at 0.16, 0.32, 0.43, and 0.61 Hz (maximum current limited to ±5 mA) which has been shown to reliably disrupt the vestibular system (MacDougall et al., 2006; Moore et al., 2006).

We tested an additional set of 15 participants from the York University undergraduate participant pool. One participant had to abandon the experiment because the sensations induced by GVS were too uncomfortable for them, which made for a final sample size of n = 14 participants. Data collection for this experiment occurred between May and December 2025.

The visual stimulus was the same as for Experiment 1, except participants performed four blocks of 60 trials (of around 3 minutes duration each). Each of these 60 blocks were composed of 30 1g trials and 30 -1g trials (interleaved; with initial vertical velocities, horizontal velocities and occlusion percentage randomized as for Experiment 1). The blocks alternated between dGVS and sham, and participants were counterbalanced for which block they started with. The whole experiment was performed seated for participant safety because the vestibular noise induced by GVS can lead to balance problems and unstable posture.

We used the same statistical model as for Experiment 1, except that GVS (GVS versus sham) replaced posture. In this experiment, 161 out of 3360 trials were excluded (4.8%).

Results are summarized in Figure 5. No difference contrasts were significant (see Appendix A for details).

**Figure 5.**
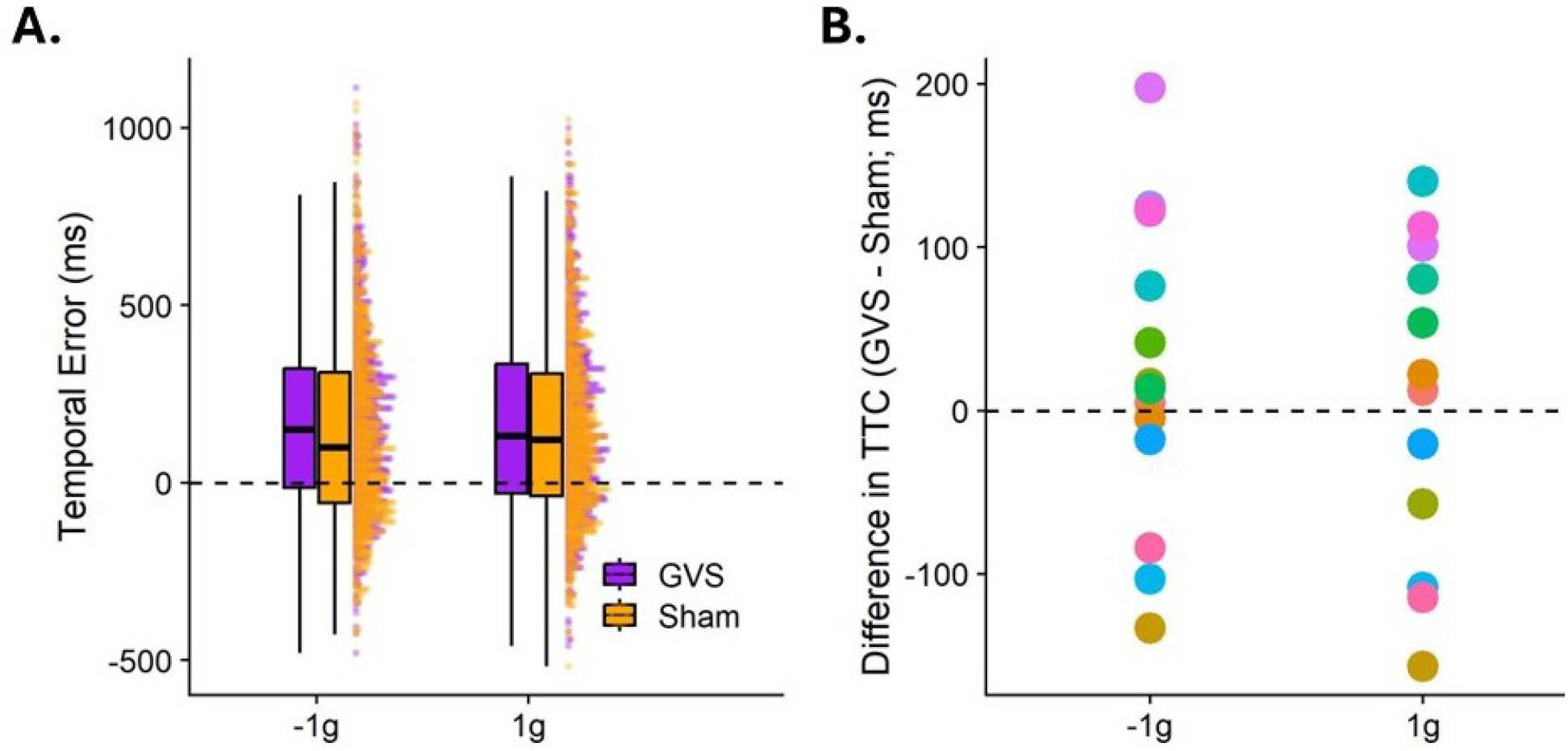
A. Full distributions of temporal errors (y axis) per gravity level (1g versus -1g, x axis) and GVS condition (GVS versus sham, color-coded). The boxplots indicate the 25^th^ and 75^th^ percentile (box), while the whiskers indicate the 2.5 times the interquartile range, and the horizontal bar in the middle corresponds to the median. The dashed line indicates perfect performance, i.e., a temporal error of 0. B. Mean difference in TTC between GVS and sham (y axis) for the two gravity conditions (x axis). Participants are color-coded. The dashed line indicates equal performance between GVS and Sham. Values above the line indicate that the participant responded later in the GVS condition than in the sham condition, and values value show that the participant responses earlier in the GVS condition than in the sham condition.

## Discussion

Contrary to our hypotheses, we found no differences between 1g and -1g, nor between sitting and lying or GVS and sham.

The absence of a difference between 1g and -1g is an intriguing finding that contrasts with a substantial body of research showing higher accuracy (less bias) when object motion is governed by Earth gravity rather than in the presence of arbitrary accelerations or an inverted Earth gravity (Bosco et al., 2015; Ceccarelli et al., 2018; Gaveau et al., 2011; Indovina et al., 2005; Jörges et al., 2018; Jörges & López-Moliner, 2017, 2019, 2020; Zago et al., 2004b, 2004a, 2008, 2009, 2010, 2011a). This was especially unexpected in the standing upright position, where the gravity prior and vestibular input are in agreement and we would expect the perceptual system to be able to use the additional information provided to better estimate time-to-contact. One potential explanation might be that our participants did not interpret our stimulus as real enough, which therefore failed to activate their internal representation of Earth gravity. It is also possible that there were not enough visual polarity cues in the simulation to indicate the local direction of simulated gravity. While this seems somewhat unlikely considering that some of the studies cited above involve much less realistic 2D stimuli, it remains our best explanation given the overwhelming evidence available in the literature.

## Open Science

The programs to present stimulus, the Unity project used to generate the programs, as well as all data and the code for all analyses are available for download at Open Science Foundation (https://osf.io/vf7qm/).

## Acknowledgements

BJJ and LRH co-designed the experiment. BJJ programmed the stimulus, analyzed and interpreted the data and wrote the manuscript, on which LRH provided extensive feedback. We thank Ahmed Nadeem and Fatemeh Ghasemi for their help with data collection. We further thank the Canadian Space Agency for partial funding of this research. BJJ was funded through a postdoctoral grant from York University’s Center for Vision Research, while LRH is holder of a grant from the Natural Sciences and Engineering Research Council of Canada.

## Appendix A Overview of All Difference Contrasts

**Table 1.** All difference contrasts for all independent variables in the three linear mixed models fitted for the three experiments, along with the 98.3% confidence intervals and an indicator whether the confidence interval includes zero (“significance”). All values in ms.

|  | Difference Contrast | 98.6% CI (Lower end) | CI_98.6% CI (Upper end) | Significance |
| --- | --- | --- | --- | --- |
| **Experiment 1: Standing vs Lying Supine** |  |  |  |  |
| Intercept | 170 | 60 | 265 | * |
| Posture (Standing vs Supine) | -20 | -104 | 51 | n.s. |
| Gravity (1g vs -1g) | 77 | -61 | 209 | n.s. |
| Interaction | -8 | -26 | 11 | n.s. |
| Vertical Velocity | 7 | -4 | 18 | n.s. |
| Horizontal Velocity | 0 | -2 | 2 | n.s. |
| **Control Experiment: Lying Supine (vs Standing) – Reaction Times** |  |  |  |  |
| Intercept | -53 | -100 | -1 | * |
| Posture (Standing vs Supine) | -1 | -18 | 19 | n.s. |
| Gravity (1g vs -1g) | -2 | -85 | 84 | n.s. |
| Interaction | 1 | -11 | 13 | n.s. |
| Vertical Velocity | 0 | -7 | 7 | n.s. |
| Horizontal Velocity | 0 | -1 | 1 | n.s. |
| **Experiment 2:**<br>**GVS vs Sham** |  |  |  |  |
| Intercept | 159 | -16 | 335 | n.s. |
| Sham (vs. GVS) | -28 | -87 | 40 | n.s. |
| Gravity (1g vs -1g) | 62 | -133 | 286 | n.s. |
| Interaction | 6 | -19 | 33 | n.s. |
| Vertical Velocity | 6 | -10 | 24 | n.s. |
| Horizontal Velocity | 1 | -2 | 5 | n.s. |

